# Physicochemical amino acid properties better describe substitution rates in large populations

**DOI:** 10.1101/378893

**Authors:** Claudia C. Weber, Simon Whelan

## Abstract

Substitutions between chemically distant amino acids are known to occur less frequently than those between more similar amino acids. This knowledge, however, is not reflected in most codon substitution models, which treat all non-synonymous changes as if they were equivalent in terms of impact on the protein. A variety of methods for integrating chemical distances into models have been proposed, with a common approach being to divide substitutions into radical or conservative categories. Nevertheless, it remains unclear whether the resulting models describe sequence evolution better than their simpler counterparts.

We propose a parametric codon model that distinguishes between radical and conservative substitutions, allowing us to assess if radical substitutions are preferentially removed by selection. Applying our new model to a range of phylogenomic data, we find differentiating between radical and conservative substitutions provides significantly better fit for large populations, but see no equivalent improvement for smaller populations. Comparing codon- and amino acid models using these same data shows that alignments from large populations tend to select phylogenetic models containing information about amino acid exchangeabilities, whereas the structure of the genetic code is more important for smaller populations.

Our results suggest selection against radical substitutions is, on average, more pronounced in large populations than smaller ones. The reduced observable effect of selection in smaller populations may be due to stronger genetic drift making it more challenging to detect preferences. Our results imply an important connection between the life history of a phylogenetic group and the model that best describes its evolution.

## Introduction

Quantifying the impact of natural selection on proteins is of broad interest in evolutionary biology, providing insight into the structural and functional constraints acting on proteins and how they adapt to an organism’s environment. The most widely used method for studying selection using multiple sequence alignments of protein-coding sequences is to consider the ratio of the non-synonymous substitution rate (*dN*) to the synonymous substitution rate (*dS*), often referred to as *ω* = *dN*/*dS*. These codon-based models assume that *dS* reflects the neutral rate of evolution and *dN* represents the rate after selection has acted. The *ω* measure is well-characterised, and established methods allow it to vary along a sequence, through an evolutionary tree, or a combination of both (Goldman and Yang, 1994; Muse and Gaut, 1994; Yang, 1998; Yang and Nielsen, 1998, 2002). Although useful, *ω* is coarse-grained because *dN* takes on the same value regardless of the amino acids being substituted despite widespread evidence that some amino acids substitutions are more likely to occur than others.

Early research showed that amino acid substitutions between amino acids with very different physicochemical properties occur less frequently than substitutions between more similar amino acids (Epstein, 1967), leading to the introduction of a range of scales measuring physicochemical distances (Grantham, 1974; Miyata *et al*., 1979). The original codon models used a *dN*/*dS* measure that incorporated these distances (Goldman and Yang, 1994), based on the rationale that selection against more similar amino acid substitutions ought to be weaker than against more distant ones. Subsequent research, however, found that this model frequently provided a poorer fit than the simpler M0 model, which estimates *dN*/*dS* but does not capture differences in the selective pressures acting on different amino acid substitutions (Yang et *al.,* 1998).

Other studies proposed classifying amino acid substitutions into conservative and radical substitutions, describing substitutions between similar and dissimilar amino acids, respectively. The relative amount of radical and conservative change can be described by the ratio *R/C*, which is interpreted as a measure of selective pressure acting on the two different classes of non-synonymous mutations. Several approaches have been proposed to estimate *R/C* by counting amino acid changes (Popadin *et al*., 2007; Smith, 2003; Weber *et al*., 2014; Zhang, 2000), and results suggest that radical mutations are more likely to be selected against than conservative mutations (Smith, 2003). Further, organisms with a small effective population size tend to accumulate more radical substitutions than those with a larger effective population size and more efficient natural selection (Eyre-Walker *et al*., 2002; Popadin *et al*., 2007; Smith, 2003), although see (Figuet *et al*., 2016). However, count-based methods may be unreliable in the presence of mutation- and composition biases (Dagan *et al*., 2002; Smith, 2003), given their implicit incorporation of parsimony-type scoring. Attempts to incorporate *R/C* into a more statistically justified and robust codon substitution model have so far been limited to surveys of positive selection in genes already known to contain sites with accelerated non-synonymous substitution rates (Sainudiin *et al*., 2005). Therefore, little is known about whether a model measuring *R/C* provides a significant improvement in model fit over M0, and to what extent statistical fit supports the notion that negative selection treats radical substitutions differently than conservative ones.

An alternative approach to describe variation in the selective pressures acting on amino acids is to model them at the amino acid-as opposed to the codon level. Here, “exchangeability” parameters, estimated from databases of sequences, capture the rate of change between pairs of amino acids and reflect a hazy combination of the genetic code and the selective pressures acting on the substitution. These models range from simple approaches that apply an averaged substitution process across the entire protein to sophisticated mixture models that have different substitution patterns for different sites (Jones *et al*., 1994; Le and Gascuel, 2008; Whelan and Goldman, 2001). They have been successfully used for a range of phylogenetic problems, with the mixture models proving valuable at resolving deep phylogenies (Lartillot *et al*., 2007). Modelling approaches that link the biophysical attributes of amino acid substitution models to codon models have also been developed (Seo and Kishino, 2009; Yang *et al*., 1998), but are currently not widely used.

Recently, statistical model selection methods for choosing between codon and amino acid substitution models have been devised, allowing the state space of the model that best describes a given sequence alignment to be determined (Seo and Kishino, 2009; Whelan *et al*., 2015). Analyses of collections of sequence alignments show that preference for nucleotide, amino acid or codon models is data set dependent. Moreover, the best-fitting model class is correlated with the intensity of selection as estimated by *dN*/*dS*, with more constrained alignments tending to select amino acid models and less constrained alignments selecting codon models. These results imply that the factors driving substitution rates between amino acids, and therefore differential model selection, could be driven by the selective pressures acting on biophysical properties, the structure of the genetic code, or a combination of the two (Whelan *et al*., 2015).

In light of the availability of vast quantities of sequence data spanning an enormous range of taxa, we return to the question of whether chemical distances predict selective preferences in protein-coding sequences. Previous analyses were restricted to a handful of mammalian sequences and therefore may not present a complete picture. To examine the relative selective pressures acting on different types of amino acid substitutions, we propose a codon model that separates non-synonymous substitutions into conservative or radical categories. Our approach allows *R/C* to be robustly estimated in the presence of mutational and compositional bias and long branches with multiple hits. Using a mutation-selection model we then explore how *R/C* will respond to variation in evolutionary variables, including effective population size (Ne), assuming that radical substitutions have larger negative effects on fitness than do conservative substitutions. We next assess whether the *R/C* ratios observed in empirical sequence data are consistent with radical substitutions being more disruptive and selectively unfavourable, and find that this is the case for a subset of alignments overwhelmingly belonging to taxa with large effective population sizes and strong selection. Moreover, we see a preference for amino acid models over standard codon models in the same taxa, while small population size predicts a preference for codon models. Taken together, our results confirm that chemical amino acid distance is a predictor of the strength of selection and that organismal life-history influences model selection.

## Results

### The <u>Co</u>nservative or <u>Ra</u>dical (CoRa) substitution model

The base model for describing codon substitutions is the M0 model (Goldman and Yang, 1994), which captures the relative selective pressures acting on non-synonymous substitutions through the *ω* = *dN*/*dS* parameter. Here, we propose the CoRa model, a generalisation of M0 that separates *ω* into two parameters *ω_C_* and *ω_R_,* which capture conservative and radical non-synonymous substitutions, respectively (see Materials and Methods).

The CoRa model adds one additional degree of freedom compared to M0, allowing a significance test of the relative fit of the two models by comparing the standard likelihood ratio test statistic to a 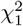 distribution. In this case M0 is the simpler model and is nested within CoRa under the condition *ω_C_* = *ω_R_*, which places conservative and radical mutations under the same amount of selective pressure. This restriction is absent in CoRa, so a measure of *R/C* describes the relative probability of fixing a radical versus a conservative mutation.

### Factors affecting R/C estimates

In the Methods section we describe two broad types of method for inferring *R/C*. The first relies on the formulation of the CoRa model and directly estimates the relative instantaneous substitution rates of *dR*/*dS* = *ω_R_* and *dC*/*dS* = *ω_C_*. The second uses observed sequences to count the relative frequency of radical to conservative substitutions, *K_r_*/*K_c_*, either: i) by using a parsimony approach to count the observable differences between sequences to obtain 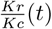, which misses cases where more than one substitution has occurred at a site; or ii) through stochastic mapping (Minin and Suchard, 2008) to obtain 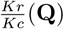, which can count all the substitutions that occurred. Using an analytic approach (see Methods) we examine how evolutionary forces can affect these different *R/C* measures, demonstrating the robustness of the *dR*/*dC* = *ω_R_*/*ω_C_* measure and the difficulties of working with *K_r_*/*K_c_* measures. All of the results presented here depend both on the classification of radical and conservative amino acid substitutions used by the CoRa model, and the structure of the genetic code, which determines the relative rates of different non-synonymous and synonymous substitutions.

### The effects of genomic variation on R/C measures

We start by assessing how *Kr*/*Kc* measures are affected by variation in the parameters of the M0 substitution model, which has equal rates of substitution between conservative and radical amino acids. Figures 1(a)-(c) show the effect of variation in *ω*, GC-content and transition-transversion (*κ*) bias on the different *Kr*/*Kc* measures. The first observation for all figures is that *Kr*/*Kc* is not equal to one even when there is no difference between conservative and radical substitution rates. The deviation from one matches previous observations (Eyre-Walker *et al*., 2002; Weber *et al*., 2014) and arises because radical substitutions are more likely to occur by chance due to the structure of the genetic code and the relative frequencies of the codons. Inferring the relative strength of selection acting on radical and conservative substitutions is therefore difficult when only raw *Kr*/*Kc* is considered, although changes in *Kr*/*Kc* might still be informative of a shift in those values (see below). This is not a problem for the *dR*/*dC* estimate since it is inferred directly from the substitution model, which accounts for the genetic code.

In Figure 1(a) we see that the ratio 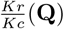 (solid black line) is not affected by changes in *ω*. This stability is expected since under the M0 model *ω_R_* = *ω_C_* = *ω*. Both *ω_R_* and *ω_C_* scale in direct proportion to *ω*, so one may expect that changes to *ω* have little effect on the *Kr*/*Kc* measure. The alternative 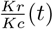 measure counts only the observable changes occurring on a branch of length *t* and is affected by variation in *ω* because its values are affected by multiple hits. For short distances where few substitutions are missed 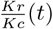 closely resembles 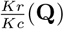. However, when *t* becomes larger there is an increasingly strong response in 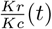 to the value of *ω* that can be attributed to the undercounting of more common non-synonymous substitutions, which becomes more severe as *ω* increases. Hence, 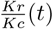 is prone to inflation for long branches. Meanwhile, variation in *ω* has no effect on the *dR*/*dC* = 1 estimate (not shown).

**FIG. 1.**
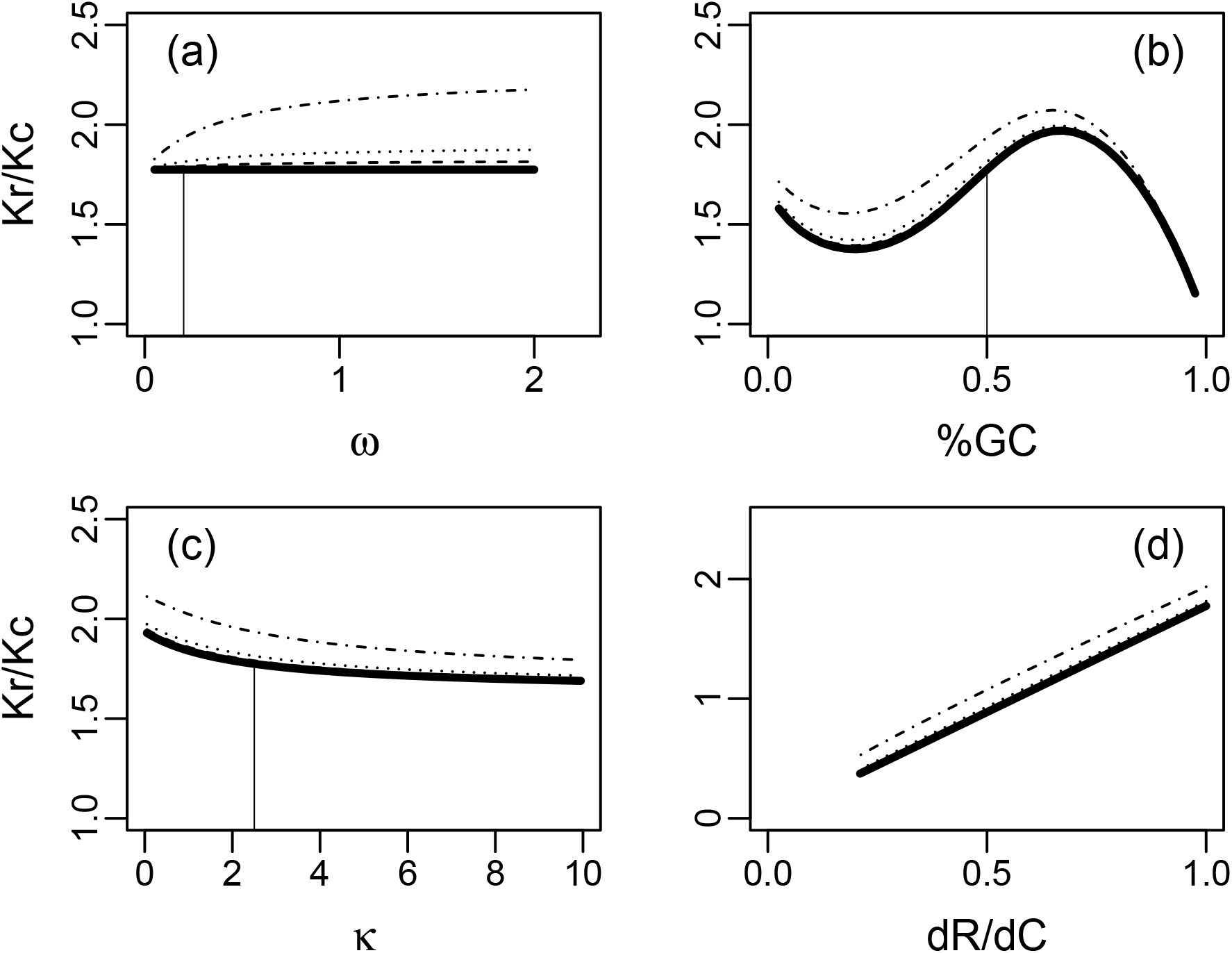
Factors affecting estimates of *K_r_*/*K_c_* obtained through counting. Solid lines represent the expected value of *K_r_*/*K_c_* when every substitution can be observed, whereas the thin dashed lines represent expected values of *K_r_*/*K_c_* observable between a pair of sequences after differing amounts of time: wide-dash for t=0.1; dotted-dash for t=0.25; and dot-dash for t=1.0. In all panels but d), *ω_R_* = *ω_C_* = *ω*. The vertical lines indicate the values of *ω*, %GC and *k* held constant across the other panels.

The effect of %GC-content and transition-transversion rate on *R/C* is shown in Figure 1(b) and 1(c), respectively. For simplicity, the effect of %GC is assessed by changing the GC content of each codon position in an identical manner to obtain a given %GC value. This approach is simpler than what occurs in nature, where GC3 is more variable than GC1 or GC2, but provides an adequate insight into the overall effect of %GC variation. Given that the CoRa model accounts explicitly for variation in %GC and *κ*, estimates of *dR*/*dC* are unaffected. In contrast %GC variation has a substantial and non-monotonic effect on both 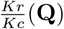 and 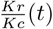. The effect of *κ* is comparatively smaller, but there is still a negative non-linear correlation between the value of *k* and 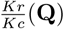 and 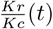. The length of time *t* tends to inflate 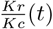 estimates in a manner dependent on both GC and *κ*, with the magnitude of the increase reflecting the specific %GC or *κ* value examined.

Finally, Figure 1(d) shows how the different *K_r_*/*K_c_* measures respond to variation in the intended target variable *dR*/*dC*. There is a clear linear relationship between *dR*/*dC* and *K_r_*/*K_c_* measures, with the slope of the line being determined by a complex function of the parameters in the M0 model under the conditions considered in the preceding panels, the genetic code, and M_rad_. This observation is encouraging since it suggests that although *Kr*/*Kc* might not be informative about the absolute selective pressures acting on conservative versus radical substitutions, relative changes in *Kr*/*Kc* might correspond to clear increases or decreases in those selective pressures. When comparing genomic sequences with the same %GC and *κ*, this relationship applies, but it is unlikely to hold for general comparisons where the values of these parameters vary.

To provide an idea of how compositional variation might affect *Kr*/*Kc* we examine two sets of M0 parameters: i) %GC = 0.5 and *κ*= 2.5 where the slope of the line is 1.77; and ii) %GC = 0.65 *κ* = 2.5 where the slope of the line increases to 1.97 (not shown in figure). The differing slope makes comparisons between regions with different properties problematic since it is difficult to disentangle changes in *Kr*/*Kc* resulting from changes in selection and those resulting from differences in sequence properties. From these results, we conclude that both *Kr*/*Kc* measures are flawed and difficult to interpret, and *dR*/*dC* hence provides the most natural and accurate estimates of the relative selective pressures acting on conservative and radical changes in proteins. We, therefore, consider only *dR*/*dC* in the following sections.

### Effective population size

The relative rates of radical to conservative substitution have been suggested to be associated with or even be suitable to predict *N_e_* (Nabholz *et al*., 2013). To explore how variation in *N_e_* relates to changes in the rates of non-synonymous substitution, we consider a simple mutation-selection model (see Materials and Methods).

Figure 2(a) shows how *ω* responds to variation in *N_e_* for various levels of negative selective pressure, *s*, where *s_c_* and *s_r_* represent selection on conservative and radical changes, respectively. The solid line represents the case where *s* = *s_c_* = *s_r_* = −0.0020 and demonstrates how *ω* changes from approximately 1.0 when *N_e_* is small to approximately 0.0 as *N_e_* becomes very large. Under the log transform of *N_e_* the different values of *s* transpose this curve on the x-axis, with the thin lines showing *s_c_* = {−0.0015,−0.0010,−0.0005} from left to right. Note that this leads to a diminishing effect of *N_e_* on *ω* as purifying selection weakens. For example, to obtain a decrease in *ω* from 0.75 to 0.25 under *s* = −0.0020 (thick line) requires an increase in *N_e_* of around 900, whereas the same reduction in *ω* for *s* = −0.0010 (dash-dot line) requires an increase in *N_e_* of around 1800.

When considering *R/C* we are interested in the ratio of differential selective pressures affecting different types of non-synonymous change, with the assumption that conservative substitutions are more likely to be fixed than radical substitutions, due to stronger selection acting against the latter (selective coefficients: *s_c_,s_r_* < 0;*s_c_*>*s_r_*; that is, *s_c_* is less negative than *s_r_*). Figure 2(b) shows how *dR*/*dC* responds to variation in *N_e_* according to equation (9) (see Materials and Methods), with each thin dotted line corresponding to different ratios of *s_r_*/*s_c_* = 4 (dotted; left), 2 (dot-dash; mid), 4/3 (dashed; right). These selective coefficients will produce a range of observed *dR*/*dC* values depending on *N_e_*. For sufficiently small populations *dR*/*dC* → 1 because selection is not strong enough to differentiate between radical and conservative mutations. Meanwhile, for sufficiently large populations *dR*/*dC* → 0 as radical substitutions are far less likely to fix than conservative substitutions. (Note that as *N_e_* gets large enough neither radical nor conservative substitutions are likely, but if one did occur it would likely be conservative under this model.)

We find that different ratios of selective coefficients for conservative and radical mutations do indeed respond differently to variation in *N_e_*, so for a fixed population size comparisons of *dR*/*dC* are informative of the selective pressures acting during evolution. For instance when comparing two genes from the same set of taxa, a lower value of *dR*/*dC* in one gene might be directly indicative of relatively stronger selection acting on radical mutations for that gene. Figure 2(b) demonstrates that these differences are difficult to interpret directly (note the non-linear spacing between *dR*/*dC* values representing different ratios of selective coefficients for a given *N_e_*). Nevertheless, relative rankings would be informative.

**FIG. 2.**
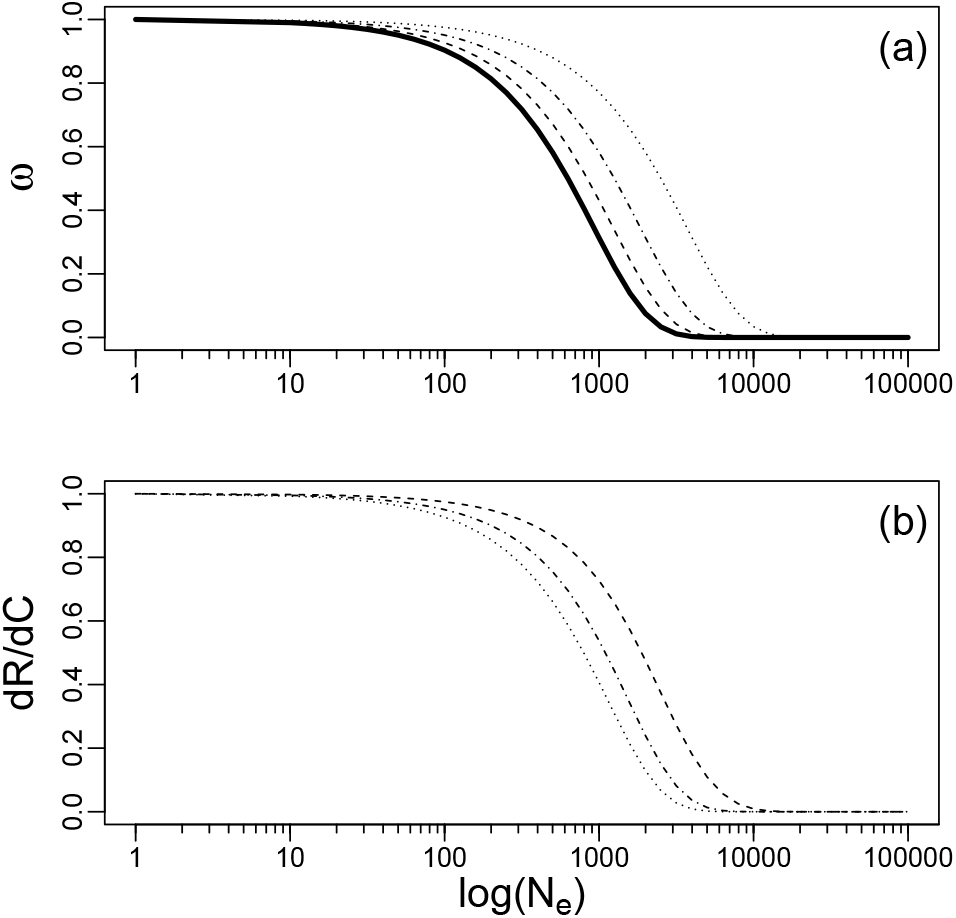
The effect of *N_e_* on (a) *ω* estimates, and (b) *dR*/*dC* estimates under the mutation-selection model. See text for more details.

Genomic analyses of amino acid substitution patterns

The previous section outlines how we expect *dR*/*dC* to respond to different values of *N_e_* and the range of values it might take. Next, we investigate whether results from real data match these expectations by examining a range of phylogenomic datasets. It is not possible to obtain accurate estimates of *N_e_* directly, so we separate our data *a priori* into two broad categories of high and low *N_e_* based on other authors’ observations. Mammals, birds and vertebrates are assumed to have relatively low *N_e_* relative to yeast, arthropods and insects, which are assumed to have relatively high *N_e_*.

### Effective population size predicts relative rates of conservative and radical substitutions

Comparisons between the CoRa and M0 models provide a direct measure of statistical support for the notion that radical and conservative substitutions are subject to different selective pressures. Figure 3 shows the distributions of the likelihood ratio test statistic, 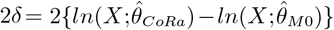, for our low and high *N_e_* genomic datasets. Here, 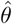 refers to the parameter estimates from the model and X to the data. We find that the distributions for 2*δ* differ markedly between the two sets, with the majority of high *N_e_* alignments falling above the 95% significance cutoff 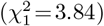, whereas a large fraction of the low *N_e_* alignments are below the threshold. That is, CoRa provides a better fit than M0 for groups with high *N_e_*, demonstrating significant support for *ω_R_* ≠ *ω_C_*, but often gives no significant improvement in fit for low *N_e_*, where we find little support for differences between the rates of radical and conservative substitutions.

**FIG. 3.**
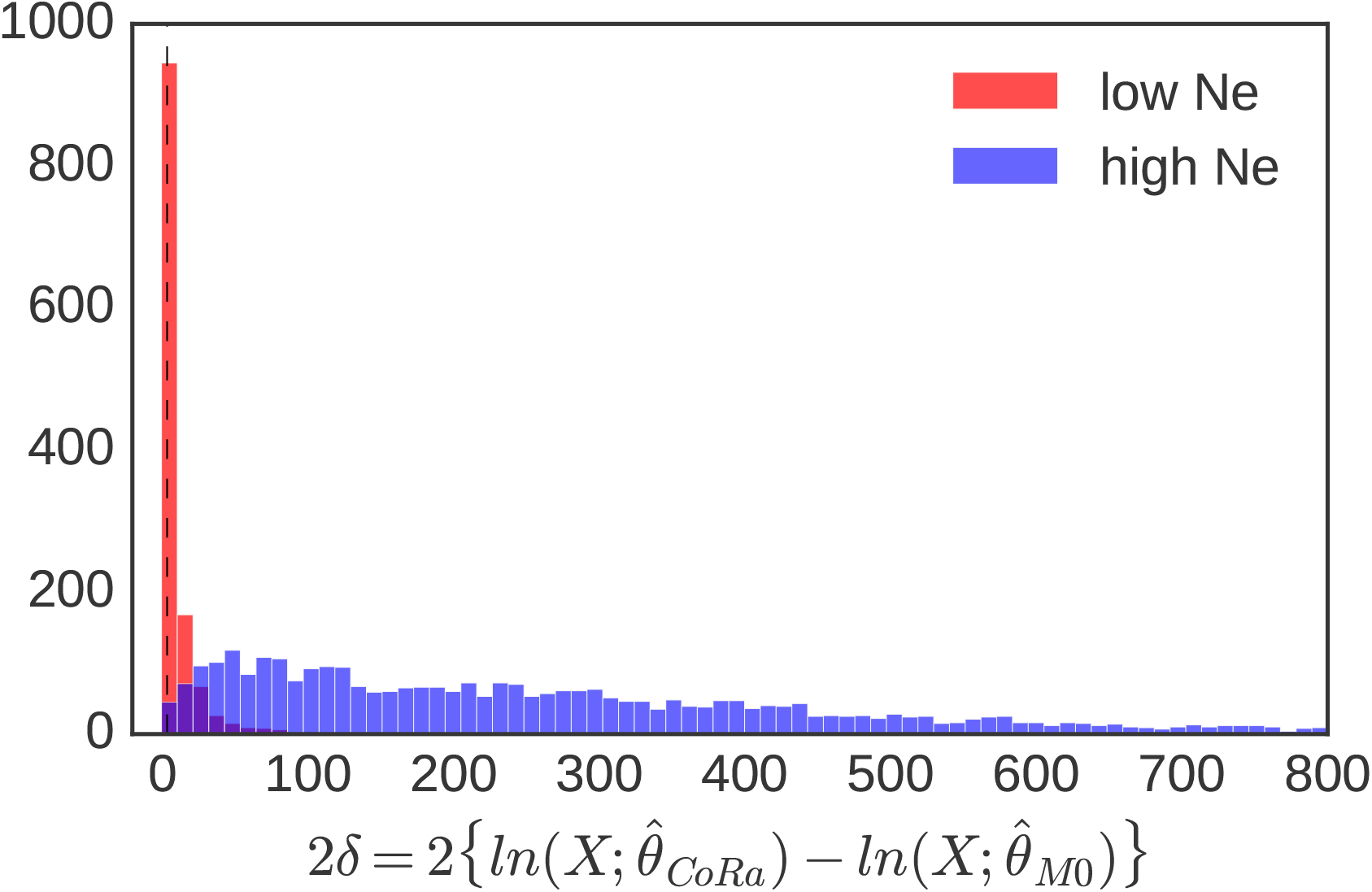
The histogram shows the distribution of the 25 statistic for CoRa compared to M0 for alignments from groups with small populations (red) compared to large populations (blue). The dashed vertical line indicates the threshold above which the statistic is significant (3.84). Significant likelihood ratio tests indicate a preference for CoRa over M0. CoRa describes alignments from large populations significantly better than M0. These results are not explained by alignment length (see Supplementary Materials, Figure S3).

There are two possible explanations for this difference in model fit. First, there might be no significant difference in the selective constraints acting on conservative (*s_c_*) and radical (*s_r_*) substitutions in the vertebrate data we examined. This explanation is implausible since there is no reason to believe that the set of proteins and their functions and levels of constraint differ between the high and low *N_e_* sets, as both categories include genome-scale data and are not restricted to curated (potentially biased) phylogenetic markers. Second, the rates of fixation of conservative and radical substitutions in low *N_e_* populations might be similar for much of the proteome resulting in *dR*/*dC* values close to 1, whereas high *N_e_* populations have more noticeable differences in *dR* and *dC*. This second explanation is consistent with predictions from Figure 2. The fact that, the whole distribution is shifted also confirms that, our observations are not. driven by any individual alignment.

To further explore this hypothesis we examine Figure 4, which plots 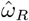 from the CoRa model against 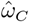. For the high *N_e_* datasets, we find the dashed blue lowess line falls below *x* = *y* suggesting 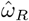 is systematically lower than 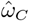. Combined with the likelihood ratio test results, this supports the hypothesis that radical substitutions are, on average, subject to stronger negative selective pressure, consistent with the expectation that they are more often deleterious. We also observe that the high *N_e_* datasets tend to have lower estimates for both *ω_C_* and *ω_R_*. In contrast, for the low *N_e_* genomic data the values of 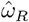 and 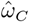 tend to be much closer to each other, with the lowess line closer to *x* = *y*, demonstrating that selection is less efficient at differentiating between radical and conservative substitutions. These observations are all consistent with our theoretical expectations and with *N_e_* leading to differences in the inferred patterns of amino acid substitution.

**FIG. 4.**
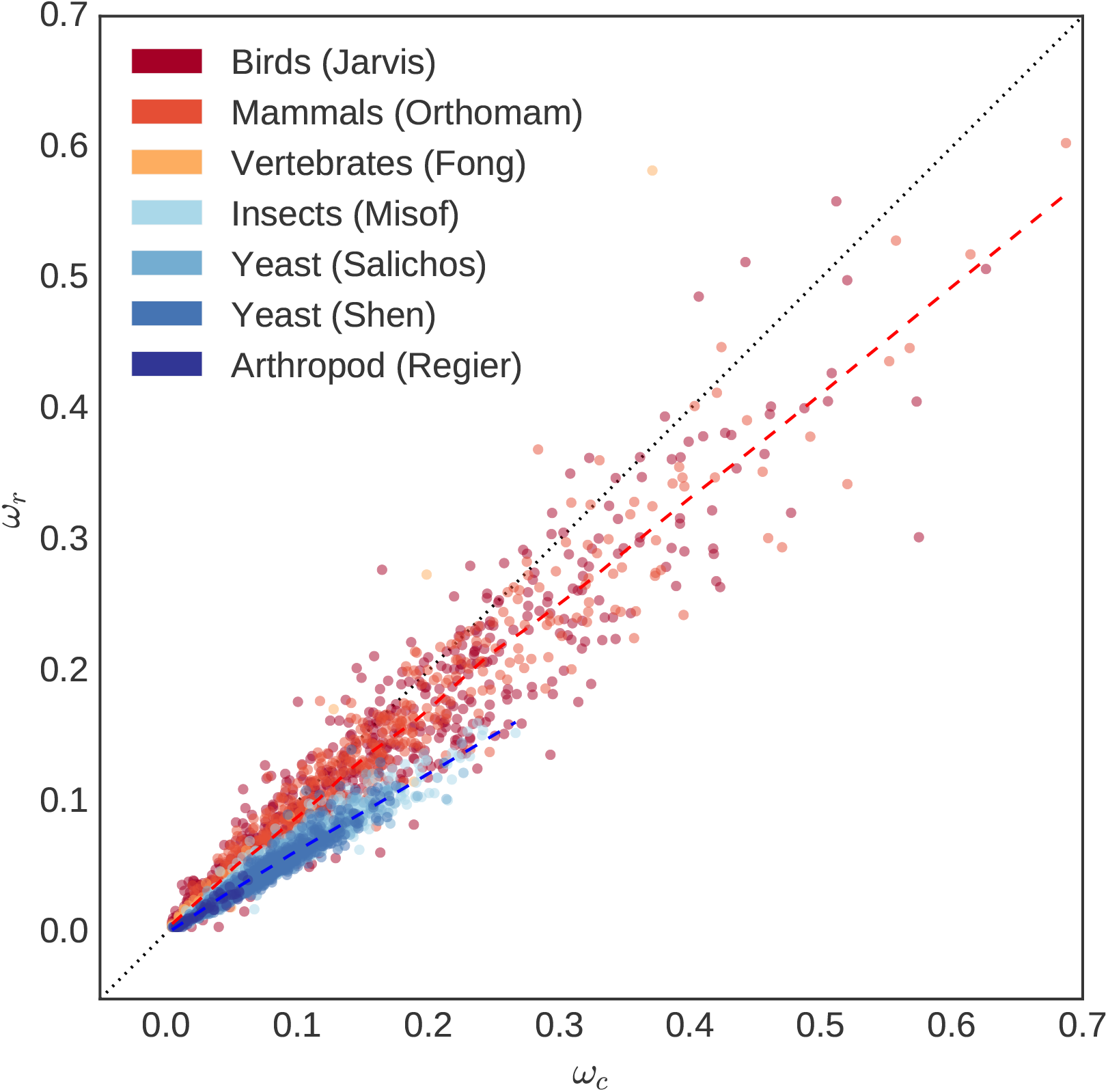
Phylogenomic datasets from taxa with different population sizes differ in terms of the relationship between 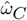 and 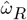. The blue and red points represent low *N_e_* and high *N_e_* phylogenomic datasets, respectively. Each point represents an individual gene and its shade indicates the phylogenomic data set it is drawn from. Although 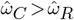 for the majority of sequences from large *N_e_* groups (insects, arthropods, yeast), vertebrate sequences show a mixed pattern and overall higher rates of radical substitution. The dotted black line indicates x = y, that is, 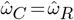. The blue lowess line represents putatively high *N_e_* groups, while red represents low *N_e_* groups.

### Effective population size drives model state-space selection

An alternative approach to incorporating the selective pressures acting on amino acid changes with different effects on biophysical properties is to describe the substitution process at the amino acid sequence level. Performing phylogenetic model selection across state-spaces using the approach of Whelan *et al*. (2015) allows us to directly compare M0-based codon models, which do not account for variation in selection acting on amino acid substitutions, to amino acid models, which include empirically derived exchangeability parameters capturing variation in rates of substitution between amino acids. Following our observation that *N_e_* affects radical and conservative substitution rates, we might expect high- and low *N_e_* populations to systematically prefer different types of model. We used the ModelOMatic program to infer both the best fitting model and the state-space of the preferred model.

Table 1 shows the overall preferred state-space for each of our phylogenomic datasets. Our inferences are consistent with both our theoretical observations and our CoRa-based analyses: high *N_e_* datasets overwhelmingly select amino acid models, which account for biophysical differences between amino acids. In contrast, low N_e_ datasets select codon models, which do not (see Table 1). Hence, models that incorporate information about amino acid properties, either in the form of binary classification via M_rad_ or a matrix describing the probability of exchange, provide a better fit for sequences from large populations with more effective selection. Here, the varying probabilities of exchange between amino acid pairs described by empirical models are analogous to assigning different selective coefficients to different classes of substitution in the CoRa model (Fig 2a).

**Table 1.** Model selection results by alignment data set. CoRa is selected when the likelihood ratio test statistic 25 is significant.

| Group | No of species | State space of preferred model | CoRa selected and $\omega_R < \omega_C$ | $\omega$ | Pop. size |
| --- | --- | --- | --- | --- | --- |
| Mammals (Douzery *et al.*, 2014) | 43 | Codon (100%) | 202/473 (43%) | 0.1152 | Small |
| Birds (Jarvis *et al.*, 2015) | 48 | Codon (100%) | 276/725 (38%) | 0.1027 | Small |
| Vertebrates (Fong *et al.*, 2012) | 129 | Codon (100%) | 17/66 (26%) | 0.0213 | Small |
| Yeast (Salichos and Rokas, 2013) | 23 | AA (97%) | 665/669 (>99 %) | 0.0615 | Large |
| Yeast (Shen *et al.*, 2016) | 96 | AA (97%) | 1214/1217 (>99 %) | 0.0518 | Large |
| Arthropods (Regier *et al.*, 2010) | 75 | AA (98%) | 67/68 (99 %) | 0.0164 | Large |
| Insects (Misof *et al.*, 2014) | 103 | AA (96%) | 1380/1389 (>99 %) | 0.0533 | Large |

To further explore these differences in preferred state-space we examine the difference in AIC between the best-fitting codon model and the best-fitting amino acid model for each alignment. Figure 5 shows the low *N_e_* datasets (red) universally and strongly select codon models. In contrast high *N_e_* datasets (blue) have a lower level of support for selecting amino acid models, which nevertheless encompasses the vast majority of alignments. The relatively weaker support for amino acid models may reflect their empirical derivation, their lower parameterisation, and their grouping codons together when describing amino acids. That is, they discard information about the evolution of individual sequences that codon models are able to capture. They might, therefore, be less well tuned to the fine-grained process, but their ability to describe average fitness differences between amino acids overcomes that limitation.

**FIG. 5.**
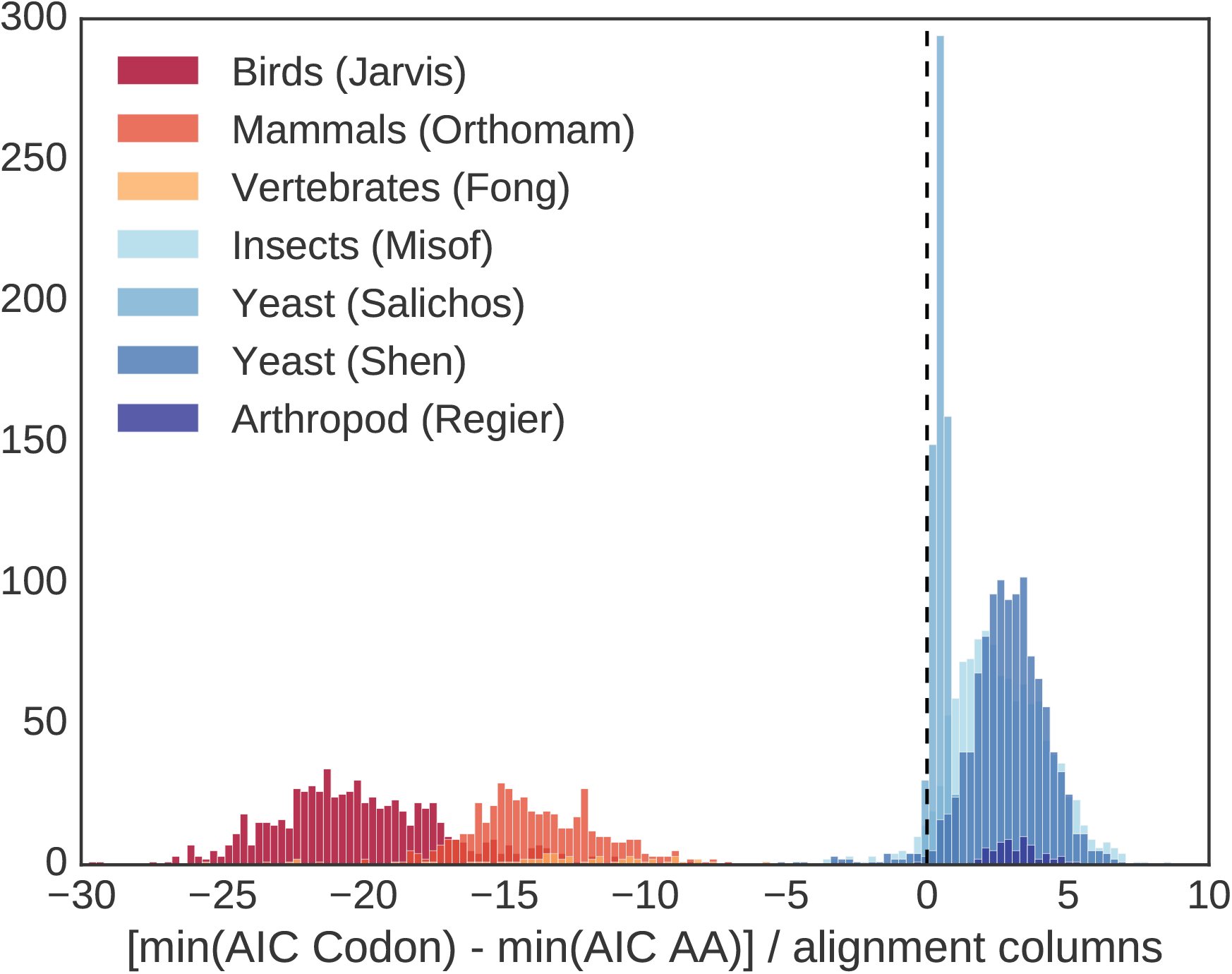
Density plot showing ΔAIC for the best-fitting AA model versus the best-fitting codon model. Blue shades indicate high Ne, while red shades low Ne. Each data point represents one alignment.

We have also considered the possibility that the amount of divergence affects our ability to detect preferences for different types of amino acid substitutions. We find that the tree lengths (sum of branch lengths) from the low *N_e_* datasets tend to be substantially shorter than those from the high *N_e_* datasets, but this difference does not seem adequate to account for the differences in the substitution process between the two groups. When examining genes with similar tree lengths across datasets the selection of CoRa versus M0 or amino acid versus codon model appears to be primarily predicted by whether the gene has evolved under high or low *N_e_*. See Supplementary Material for more information.

## Discussion

In this study, we examine how the physicochemical properties of amino acids might improve the ability of phylogenetic models to describe protein evolution. We predict a direct link between rates of amino acid substitution and effective population size (*N_e_*) mediated by variation in the efficacy of natural selection. One aspect of amino acid substitution patterns that has recently received renewed attention is whether more chemically distant changes are less frequently fixed (Figuet *et al*., 2016; Nabholz *et al*., 2013; Weber *et al*., 2014). We show that fully investigating this relationship requires a parametric substitution model since simplified counting methods are susceptible to a range of established biases, such as variation in GC content and transition-transversion bias (Dagan *et al*., 2002; Smith, 2003), which can make the results challenging to interpret. Additionally, we find that long branch lengths and high proportions of non-synonymous substitutions can exacerbate these issues. We therefore propose the <u>Co</u>nservative-<u>Ra</u>dical (CoRa) codon model that allows different types of amino acid substitution to be treated separately.

We test our theoretical predictions by comparing the relative statistical fit of phylogenetic substitution models across two broad bins of population sizes. We find that phylogenetic data from low *N_e_* populations provides little information to differentiate between broad classes of amino acid change, and simple models in the codon state-space tend to fit better than more general models in the amino acid state-space. In contrast, for high *N_e_* populations we observe distinct differences between rates of amino acid substitution and complex models in the amino acid state-space fit substantially better than simple codon-based models. In the latter case, we also confirm that radical substitutions are more strongly selected against than conservative substitutions, consistent with their being more deleterious. These results hold across several different phylogenetic datasets and, to the best of our knowledge, are not accounted for by other factors, such as sequence divergence, differences in protein function between datasets or artifacts caused by low-quality sequence data.

Given that one might naively expect constraints on structure to be better captured when amino acid preferences are considered, it may appear counterintuitive that some sequences are better described by simpler models. One explanation is that the structure of the genetic code leads to an inherent bias towards less disruptive changes compared to what would be observed under a random alternative code (Freeland and Hurst, 1998; Haig and Hurst, 1991). The genetic code alone is unlikely to explain, however, the preference for simple models in low *N_e_* populations. It is more likely that we lack either the statistical power or model resolution to detect that selective differences between broadly binned amino acids for low *N_e_* populations. More phylogenetic diversity or longer sequences might help with statistical power, and methods that allow across-site variation in the non-synonymous rates might also be helpful.

The very clear support for codon-over amino acid models observed for low *N_e_* alignments may also reflect the drawbacks of not considering information about the codon-level processes that influenced the evolution of the sequences (Seo and Kishino, 2008). The observation that a subset of the same sequence alignments that reject amino acid models nonetheless select CoRa over M0 aligns with the idea that accounting for the structure of the genetic code does offer advantages. Overall, our results are in accord with the notion that selective constraints on amino acid changes are, on average, greater than the relative constraints on radical changes compared to conservative ones (Smith, 2003) and that this is more readily detectable in high *N_e_* populations.

That the rate of molecular evolution correlates with population size as a function of the strength of selection is not surprising and has previously been widely discussed (Bromham, 2011; Hua and Bromham, 2017; Woolfit, 2009). Here, we confirm that this observation also extends to the ratio of radical to conservative substitutions, *R/C*, in accord with theoretical expectations. Although this relationship has previously been hinted at, we can now rule out the possibility that it is merely an artifact of the effect of compositional bias on count-based estimates of *R/C* (see discussion in Weber *et al*. (2014)). In addition, we find that life history does not just affect rates of evolution but also more broadly predicts which models are suitable for describing them.

Our results may help explain many previous observations in the literature and provide insight into possible causes for the differences in relative fit of existing phylogenetic models. There is a wide variety of available amino acid substitution models, trained on data ranging from a handful of mammalian mitochondrial sequences (Yang *et al*., 1998) to vast groups of species covering the entire tree of life (Le and Gascuel, 2008). The difference between these models is usually discussed in terms of the varying protein contents they describe (Keane *et al*., 2006) or the parameters they infer, but our results suggest that other factors such as differences in *N_e_* affecting their training data could also play an important role. Our results suggest, for instance, that amino acid substitution models models trained on low *N_e_* mammalian data might be more reflective of the genetic code whereas models trained across the tree of life, which consider sequences from extremely high *N_e_* species, could capture more subtle preferences driven by physicochemical differences between amino acids.

The effect of *N_e_* on substitution rates may also explain the limited improvement in fit observed in early attempts to incorporate physicochemical distances between amino acids into codon models (Yang *et al*., 1998). The sequences examined in these studies came from mammals, which have small populations and would therefore likely require a substantial amount of phylogenetic information to discriminate between different categories of amino acids.

Variation in *N_e_* across the tree of life and its differential impact on radical and conservative substitutions might also help explain other widely discussed observations in evolutionary biology. One example is the pervasive temporal heterogeneity in substitution rates throughout the tree of life. This heterogeneity is known to cause significant problems for phylogenetic inference (Galtier, 2001; Huelsenbeck, 2002; Tuffley and Steel, 1998; Wang *et al*., 2006; Whelan, 2008; Whelan *et al*., 2010), although its exact causes remain nebulous. Our results suggest that variation in *N_e_* across the tree might be an important causal factor for this temporal heterogeneity. For instance, a change from a low to high *N_e_* along a branch might lead to a switch from ‘on’ to ‘off’ in a covarion process (Tuffley and Steel, 1998). Intriguingly, our results suggest this covarion-type switch could be an extreme example and there might be more subtle, but pervasive, changes in amino acid substitution patterns across the tree, whereby radical changes are more or less efficiently purged by selection depending on *N_e_*. In addition to being a clear model violation, there might appear to be convergent evolution in taxa across the tree with high *N_e_* simply because there are fewer amino acids that are accepted by selection at individual sites on these lineages, even when there are multiple amino acids that are roughly functionally equivalent. In such cases, methods that look for convergent shifts in amino acid profiles rather than shifts towards a single amino acid (Rey *et al*., 2018) may be more robust. At the very least our argument suggests one should be cautious when building trees from groups of taxa with very different life histories since even the most sophisticated amino acid substitution models, such as CAT and PMSF (Blanquart and Lartillot, 2008; Lartillot and Philippe, 2004; Si Quang *et al*., 2008; Wang *et al*., 2018) only account for spatial heterogeneity along the sequence and cannot capture variation in substitution patterns through the tree.

Finally, our findings also suggest that one path towards capturing and understanding protein evolution is to begin explicitly incorporating *N_e_* into phylogenetic models. One particularly promising route would be the implementation of mutation-selection models (Halpern and Bruno, 1998; Tamuri *et al*., 2012; Thorne *et al*., 2012; Yang and Nielsen, 2008). These MutSel models attempt to disentangle the effects of mutation and selection, allowing the product *N_e_* * *s* to be estimated for different types of substitutions from phylogenetic data. The models might then be extended to estimate relative values of *N_e_* for branches or clades. While the results we present in this manuscript allow us to establish that radical substitutions are preferentially selected against, a MutSel model might also allow us to obtain a more detailed picture of what constitutes a “radical” substitution in a specific site or protein. An alternative approach might be to use MCMC to estimate temporal mixture models with different substitution models at different sites on different branches. Either way, unlocking the relationship between *N_e_* and substitution rate may help resolve some of the most difficult questions in phylogenetics.

## Methods

### Models

#### Existing codon models

Standard phylogenetic methods for analysing 735 codon sequences are typically based around the M0 model derived from Yang and Goldman (Goldman and Yang, 1994). This substitution model is defined through the off-diagonal elements of the instantaneous rate matrix, where *ω* = 740 *dN*/*dS*, as follows:

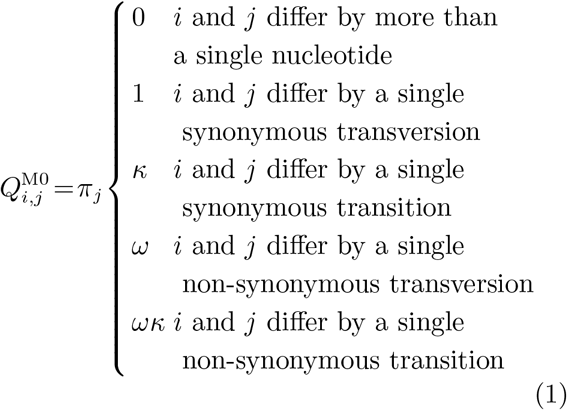

Following standard approaches, a substitution model can be used to produce transition matrices describing the probability of change between different codons over time *t*, such that *P*^M0^(*t*) = *e*^Q*t*^. The diagonal elements of the instantaneous matrix are set to the negative row sums to ensure the models produce a valid set of probability matrices for different time points. This model assumes that all amino acid mutations are equally likely to be fixed, which ignores differences in the physicochemical properties of the amino acids.

### The CoRa codon model describing radical and conservative amino acid change

The frequency at which conservative and radical amino acid substitutions occur provides some insight into the evolutionary forces acting on proteins (Sainudiin *et al*., 2005; Smith, 2003; Zhang, 2000), but there is limited discussion regarding how different estimates of these values relate to the well-studied parameters of the M0 model. Here we present the Conservative and Radical (CoRa) codon model of amino acid substitution, which divides *ω* = *dN*/*dS* from M0 into to the two subcategories *ω*_C_ = *dC*/*dS* and *ω*_R_ = *dR*/*dS* in the following instantaneous rate matrix:

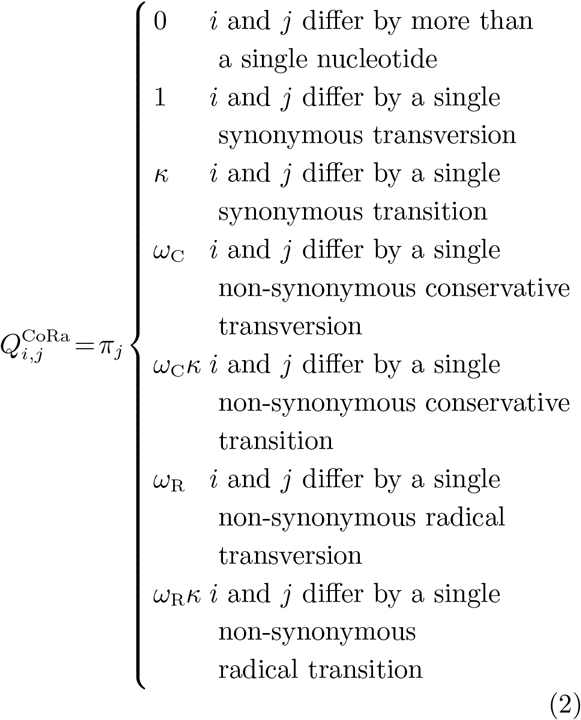

In order to fully resolve this matrix we need to define the radical matrix M_rad_, which is a 20×20 binary matrix that labels each amino acid substitution *i* → *j* as a 0 or 1, representing a conservative and a radical amino acid substitution, respectively. Here, we distinguish between polar and non-polar amino acids and amino acids with large and small volumes, where Y, W, H, K, R, E, Q, T, D, N, S and C are polar and L, I, F, M, Y, W, H, K, R, E and Q are large. Any substitution involving a change in category for polarity or volume is considered radical, whereas substitutions altering neither property’s category are considered conservative (Sainudiin *et al*., 2005). In a preliminary study, we examined other conservative and radical classifications, but the above definition tended to provide better fitting models.

Under this model we can calculate *dR*/*dC* = *ω*_R_/*ω*_C_, which is a *R/C* measure describing the relative rates of conservative and radical substitutions. Under the constraint *ω*_R_ = *ω*_C_ conservative and radical changes cannot be distinguished by selection and CoRa reduces to the simpler model M0. This nesting means that standard likelihood ratio tests with one degree of freedom may be used to assess the fit of CoRa relative to M0. When CoRa provides no significant improvement on M0 there is no evidence for differences in the rate of conservative and radical substitutions. With a significant improvement in fit, then *dR*/*dC* < 1 represents our expectation that radical changes, which are likely to disrupt the protein structure, occur at a lower instantaneous rate than conservative changes, which are more likely to maintain the current structure. A significant improvement in fit coupled with values of *dR*/*dC* > 1 represent the case where radical substitutions occur significantly more frequently than conservative substitutions.

### Counting-based K_r_/K_c_ measures and their relationship to the CoRa model

The definition of *dR* and *dC* described above has a clear relationship to the commonly used M0 model and the well defined *ω* parameter. Other studies have developed alternative *R/C* measures. Here, we therefore characterise some of these measures and show that *dR*/*dC* provides the most convenient and interpretable value. The two alternatives we examine involve different approaches to counting conservative (*Kc*) and radical (*Kr*) non-synonymous substitutions, and using *K_r_*/*K_c_* as a measure of the relative selective pressure acting on conservative and radical substitutions. The first involves tallying the total number of substitutions that occur per unit time. Here, all cases of ‘multiple hits’ where several substitutions occur at a site and can be captured through stochastic mapping (Minin and Suchard, 2008; Nielsen and Huelsenbeck, 2002) or by counting the number of conservative and radical changes over a very short period of time. Using the CoRa model the expected number of conservative and radical substitutions per unit time can be computed as:

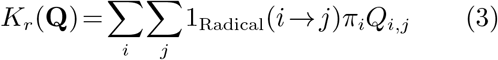

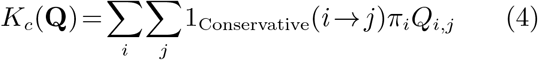

where 1_Radical_ and 1_Conservative_ are indicator functions that return 1 when *i* → *j* are radical and conservative non-synonymous substitutions, 835 respectively, and 0 otherwise. Expected values calculated using this approach will be referred to as 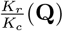 to indicate they have been derived from the instantaneous rate matrix **Q**. The second approach to estimating *K_r_*/*K_c_* is to calculate the 840 the relative numbers of conservative and radical substitutions after a given evolutionary time, *t*. This approach is essentially the same as above, but replaces **Q** with the probability matrix **P**(*t*), and fails to correct for cases where more than one 845 substitution occurs at a site. For a given value of *t* these quantities can be calculated as:

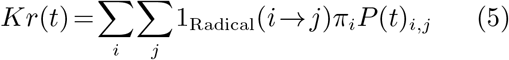

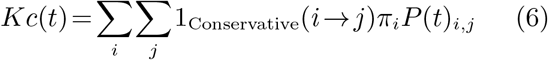

For both of these cases, the *Kr* and *Kc* measures are comparable to those obtained from amino acid sequences when counting the 850 number of radical and conservative substitutions. Estimates made using this approach will be referred to as 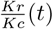.

### Defining dR/dC in terms of N_e_

Several studies have proposed that radical and conservative substitutions might predict *N_e_*, as radical substitutions are expected to be removed more effectively from large populations (Eyre-Walker *et al*., 2002). Following the formulation of mutation-selection models, where the substitution rate (*Q_ij_*) is a product of the mutation rate from *i* to *j* and the probability of fixation of *j* relative to wildtype *i*, the CoRa model describes the relative fixation probabilities through the *ω_C_* and *ω_R_* parameters. Using the weak mutation model of Golding and Felsenstein (Golding and Felsenstein, 1990) to define these parameters in terms of the selective pressures acting on conservative (*s_c_*) and radical changes (*s_r_*) for diploids we have:

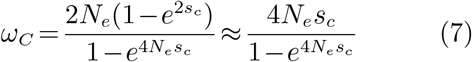

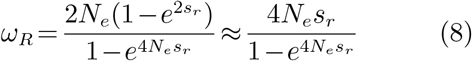

Under this approach both conservative and radical changes will be assumed to be selectively disadvantageous, which allows us to express *dR*/*dC* in terms of these selective pressures and *N_e_*:

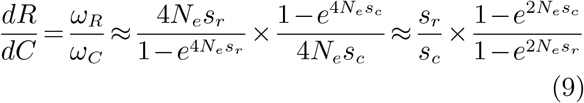

Note that this expression only allows us to examine how dR/dC varies according to *N_e_* for relative values of *s_c_* and *s_r_* since they either only occur as a fraction or in terms with *N_e_*. (The effect of *N_e_* on the values of *ω_C_* and *ω_R_* can also be computed in this manner, with Yang and Nielsen providing details (Yang and Nielsen, 2008).) Our approach allows us to analytically examine how *dN*/*dS* might respond for different values of *N_e_*, *s_c_* and *s_r_*.

### Analysis of phylogenomic datasets

We take previously published phylogenomic datasets each consisting of a set of alignments from between 23-129 OTUs from birds (Jarvis *et al*., 2014, 2015), mammals (Douzery *et al*., 2014), insects (Misof *et al*., 2014), arthropods (Regier *et al*., 2010) and yeast (Salichos and Rokas, 2013; Shen *et al*., 2016), with the range of taxonomic groups providing comparisons between a range of effective population sizes (Charlesworth, 2009; Skelly *et al*., 2009) and tree depths.

We performed basic quality control, including removing all sequences with internal stops; CTG 900 codons were masked out in the *Candida* clade due to a non-standard genetic code. To obtain codon alignments for the data of Salichos et al. (Salichos and Rokas, 2013), amino acid alignments were mapped back to nucleotide 905 sequences using PAL2NAL (Suyama *et al*., 2006).

The sequence alignments in each data set were analysed independently. The optimal state space (RY, nucleotide, codon or amino acid) for the phylogenetic analysis was determined using ModelOMatic with default settings (Whelan *et al*., 2015).

To ensure AIC values from ModelOMatic are comparable between alignments of different lengths we scaled them by the number of alignment columns present after filtering sparse columns. Purpose-written software was used to estimate parameters from M0 and the CoRa model using maximum likelihood. BioNJ trees for each multiple sequence alignment were used as a starting point to fit the models (Gascuel, 1997).

## References

Blanquart, S. and Lartillot, N. 2008. A site-and time-heterogeneous model of amino acid replacement. Molecular biology and evolution, 25(5): 842–858.

Bromham, L. 2011. The genome as a life-history character: why rate of molecular evolution varies between mammal species. Philosophical Transactions of the Royal Society of London B: Biological Sciences, 366(1577): 2503–2513.

Charlesworth, B. 2009. Effective population size and patterns of molecular evolution and variation. Nature Reviews Genetics, 10(3): 195–205.

Dagan, T., Talmor, Y., and Graur, D. 2002. Ratios of radical to conservative amino acid replacement are affected by mutational and compositional factors and may not be indicative of positive darwinian selection. Molecular biology and evolution, 19(7): 1022–1025.

Douzery, E. J. P., Scornavacca, C., Romiguier, J., Belkhir, K., Galtier, N., Delsuc, F., and Ranwez, V. 2014. OrthoMaM v8: A Database of Orthologous Exons and Coding Sequences for Comparative Genomics in Mammals. Molecular Biology and Evolution, 31(7): 1923–1928.

Epstein, C. J. 1967. Non-randomness of Amino-acid Changes in the Evolution of Homologous Proteins. Nature, 215: 355–359.

Eyre-Walker, A., Keightley, P. D., Smith, N. G. C., and Gaffney, D. 2002. Quantifying the slightly deleterious mutation model of molecular evolution. Molecular Biology and Evolution, 19(12): 2142–9.

Figuet, E., Nabholz, B., Bonneau, M., Carrio, E. M., Nadachowska-Brzyska, K., Ellegren, H., and Galtier, N. 2016. Life history traits, protein evolution, and the nearly neutral theory in amniotes. Molecular biology and evolution, 33(6): 1517–1527.

Fong, J. J., Brown, J. M., Fujita, M. K., and Boussau, B. 2012. A phylogenomic approach to vertebrate phylogeny supports a turtle-archosaur affinity and a possible paraphyletic lissamphibia. PLoS One, 7(11): e48990.

Freeland, S. J. and Hurst, L. D. 1998. The genetic code is one in a million. Journal of molecular evolution, 47(3): 238–248.

Galtier, N. 2001. Maximum-likelihood phylogenetic analysis under a covarion-like model. Molecular Biology and Evolution, 18(5): 866–873.

Gascuel, O. 1997. Bionj: an improved version of the nj algorithm based on a simple model of sequence data. Molecular biology and evolution, 14(7): 685–695.

Golding, B. and Felsenstein, J. 1990. A maximum likelihood approach to the detection of selection from a phylogeny. Journal of molecular evolution, 31(6): 511–523.

Goldman, N. and Yang, Z. 1994. A codon-based model of nucleotide substitution for protein-coding dna sequences. Molecular biology and evolution, 11(5): 725–736.

Grantham, R. 1974. Amino acid difference formula to help explain protein evolution. Science, 185: 862–864.

Haig, D. and Hurst, L. D. 1991. A quantitative measure of error minimization in the genetic code. Journal of molecular evolution, 33(5): 412–417.

Halpern, A. L. and Bruno, W. J. 1998. Evolutionary distances for protein-coding sequences: modeling site-specific residue frequencies. Molecular biology and evolution, 15(7): 910–917.

Hua, X. and Bromham, L. 2017. Darwinism for the genomic age: connecting mutation to diversification. Frontiers in genetics, 8(12).

Huelsenbeck, J. P. 2002. Testing a covariotide model of dna substitution. Molecular biology and evolution, 19(5): 698–707.

Jarvis, E. D., Mirarab, S., Aberer, A. J., Li, B., Houde, P., Li, C., Ho, S. Y. W., Faircloth, B. C., Nabholz, B., Howard, J. T., Suh, A., Weber, C. C., da Fonseca, R. R., Li, J., Zhang, F., Li, H., Zhou, L., Narula, N., Liu, L., Ganapathy, G., Boussau, B., Bayzid, M. S., Zavidovych, V., Subramanian, S., Gabaldon, T., Capella-Gutierrez, S., Huerta-Cepas, J., Rekepalli, B., Munch, K., Schierup, M., Lindow, B., Warren, W. C., Ray, D., Green, R. E., Bruford, M. W., Zhan, X., Dixon, A., Li, S., Li, N., Huang, Y., Derryberry, E. P., Bertelsen, M. F., Sheldon, F. H., Brumfield, R. T., Mello, C. V., Lovell, P. V., Wirthlin, M., Schneider, M. P. C., Prosdocimi, F., Samaniego, J. A., Velazquez, A. M. V., Alfaro-Nunez, A., Campos, P. F., Petersen, B., Sicheritz-Ponten, T., Pas, A., Bailey, T., Scofield, P., Bunce, M., Lambert, D. M., Zhou, Q., Perelman, P., Driskell, A. C., Shapiro, B., Xiong, Z., Zeng, Y., Liu, S., Li, Z., Liu, B., Wu, K., Xiao, J., Yinqi, X., Zheng, Q., Zhang, Y., Yang, H., Wang, J., Smeds, L., Rheindt, F. E., Braun, M., Fjeldsa, J., Orlando, L., Barker, F. K., Jonsson, K. A., Johnson, W., Koepfli, K.-P., O’Brien, S., Haussler, D., Ryder, O. A., Rahbek, C., Willerslev, E., Graves, G. R., Glenn, T. C., McCormack, J., Burt, D., Ellegren, H., Alstrom, P., Edwards, S. V., Stamatakis, A., Mindell, D. P., Cracraft, J., Braun, E. L., Warnow, T., Jun, W., Gilbert, M. T. P., and Zhang, G. 2014. Whole-genome analyses resolve early branches in the tree of life of modern birds. Science, 346(6215): 1320–1331.

Jarvis, E. D., Mirarab, S., Aberer, A. J., Li, B., Houde, P., Li, C., Ho, S. Y. W., Faircloth, B. C., Nabholz, B., Howard, J. T., Suh, A., Weber, C. C., da Fonseca, R. R., Alfaro-Núñez, A., Narula, N., Liu, L., Burt, D., Ellegren, H., Edwards, S. V., Stamatakis, A., Mindell, D. P., Cracraft, J., Braun, E. L., Warnow, T., Jun, W., Gilbert, M., and Zhang, G. 2015. Phylogenomic analyses data of the avian phylogenomics project. GigaScience, 4(1): 4.

Jones, D., Taylor, W., and Thornton, J. 1994. A mutation data matrix for transmembrane proteins. FEBS letters, 339(3): 269–275.

Keane, T. M., Creevey, C. J., Pentony, M. M., Naughton, T. J., and Mclnerney, J. O. 2006. Assessment of methods for amino acid matrix selection and their use on empirical data shows that ad hoc assumptions for choice of matrix are not justified. BMC evolutionary biology, 6(1): 29.

Lartillot, N. and Philippe, H. 2004. A bayesian mixture model for across-site heterogeneities in the amino-acid replacement process. Molecular biology and evolution, 21(6): 1095–1109.

Lartillot, N., Brinkmann, H., and Philippe, H. 2007. Suppression of long-branch attraction artefacts in the animal phylogeny using a site-heterogeneous model. BMC evolutionary biology, 7(1): S4.

Le, S. Q. and Gascuel, O. 2008. An improved general amino acid replacement matrix. Molecular biology and evolution, 25(7): 1307–1320.

Minin, V. N. and Suchard, M. A. 2008. Fast, accurate and simulation-free stochastic mapping. Philosophical Transactions of the Royal Society of London B: Biological Sciences, 363(1512): 3985–3995.

Misof, B., Liu, S., Meusemann, K., Peters, R. S., Donath, A., Mayer, C., Frandsen, P. B., Ware, J., Flouri, T., Beutel, R. G., et al. 2014. Phylogenomics resolves the timing and pattern of insect evolution. Science, 346(6210): 763–767.

Miyata, T., Miyazawa, S., and Yasunaga, T. 1979. Two types of amino acid substitutions in protein evolution. Journal of Molecular Evolution, 12(3): 219–236.

Muse, S. V. and Gaut, B. S. 1994. A likelihood approach for comparing synonymous and nonsynonymous nucleotide substitution rates, with application to the chloroplast genome. Molecular biology and evolution, 11(5): 715–724.

Nabholz, B., Uwimana, N., and Lartillot, N. 2013. Reconstructing the phylogenetic history of long-term effective population size and life-history traits using patterns of amino acid replacement in mitochondrial genomes of mammals and birds. Genome Biology and Evolution, 5(7): 1273–90.

Nielsen, R. and Huelsenbeck, J. 2002. Mapping mutations on phylogenies. Systematic Biology, 51(5): 729–739.

Popadin, K. Y., Polishchuk, L. V., Mamirova, L., Knorre, D., and Gunbin, K. 2007. Accumulation of slightly deleterious mutations in mitochondrial protein-coding genes of large versus small mammals. Proceedings of the National Academy of Sciences of the United States of America, 104(33): 13390–5.

Regier, J. C., Shultz, J. W., Zwick, A., Hussey, A., Ball, B., Wetzer, R., Martin, J. W., and Cunningham, C. W. 2010. Arthropod relationships revealed by phylogenomic analysis of nuclear protein-coding sequences. Nature, 463(7284): 1079–1083.

Rey, C., Guguen, L., Smon, M., and Boussau, B. 2018. Accurate detection of convergent amino-acid evolution with pcoc. Molecular Biology and Evolution, page msy114.

Sainudiin, R., Wong, W. S. W., Yogeeswaran, K., Nasrallah, J. B., Yang, Z., and Nielsen, R. 2005. Detecting site-specific physicochemical selective pressures: applications to the Class I HLA of the human major histocompatibility complex and the SRK of the plant sporophytic self-incompatibility system. Journal of Molecular Evolution, 60(3): 315–26.

Salichos, L. and Rokas, A. 2013. Inferring ancient divergences requires genes with strong phylogenetic signals. Nature, 497(7449): 327–331.

Seo, T.-K. and Kishino, H. 2008. Synonymous substitutions substantially improve evolutionary inference from highly diverged proteins. Systematic Biology, 57(3): 367–377.

Seo, T.-K. and Kishino, H. 2009. Statistical comparison of nucleotide, amino acid, and codon substitution models for evolutionary analysis of protein-coding sequences. Systematic biology, 58(2): 199–210.

Shen, X.-X., Zhou, X., Kominek, J., Kurtzman, C. P., Hittinger, C. T., and Rokas, A. 2016. Reconstructing the backbone of the saccharomycotina yeast phylogeny using genome-scale data. G3: Genes—Genomes—Genetics, 6(12): 3927–3939.

Si Quang, L., Gascuel, O., and Lartillot, N. 2008. Empirical profile mixture models for phylogenetic reconstruction. Bioinformatics, 24(20): 2317–2323.

Skelly, D. A., Ronald, J., Connelly, C. F., and Akey, J. M. 2009. Population genomics of intron splicing in 38 saccharomyces cerevisiae genome sequences. Genome biology and evolution, 1: 466–478.

Smith, N. G. C. 2003. Are radical and conservative substitution rates useful statistics in molecular evolution? Journal of Molecular Evolution, 57(4): 467–78.

Suyama, M., Torrents, D., and Bork, P. 2006. Pal2nal: robust conversion of protein sequence alignments into the corresponding codon alignments. Nucleic acids research, 34(suppl 2): W609–W612.

Tamuri, A. U., dos Reis, M., and Goldstein, R. A. 2012. Estimating the distribution of selection coefficients from phylogenetic data using sitewise mutation-selection models. Genetics, 190(3): 1101–1115.

Thorne, J. L., Lartillot, N., Rodrigue, N., and Choi, S. C. 2012. Codon models as a vehicle for reconciling population genetics with inter-specific sequence data. pages 97–110. Oxford University Press.

Tuffley, C. and Steel, M. 1998. Modeling the covarion hypothesis of nucleotide substitution. Mathematical biosciences, 147(1): 63–91.

Wang, H.-C., Spencer, M., Susko, E., and Roger, A. J. 2006. Testing for covarion-like evolution in protein sequences. Molecular biology and evolution, 24(1): 294–305.

Wang, H.-C., Minh, B. Q., Susko, E., and Roger, A. J. 2018. Modeling site heterogeneity with posterior mean site frequency profiles accelerates accurate phylogenomic estimation. Systematic biology, 67(2): 216–235.

Weber, C. C., Nabholz, B., Romiguier, J., and Ellegren, H. 2014. Kr/Kc but not dN/dS correlates positively with body mass in birds, raising implications for inferring lineage-specific selection. Genome Biology, 15(12): 542.

Whelan, S. 2008. Spatial and temporal heterogeneity in nucleotide sequence evolution. Molecular Biology and Evolution, 25(8): 1683–1694.

Whelan, S. and Goldman, N. 2001. A general empirical model of protein evolution derived from multiple protein families using a maximum-likelihood approach. Molecular biology and evolution, 18(5): 691–699.

Whelan, S., Blackburne, B. P., and Spencer, M. 2010. Phylogenetic substitution models for detecting heterotachy during plastid evolution. Molecular biology and evolution, 28(1): 449–458.

Whelan, S., Allen, J. E., Blackburne, B. P., and Talavera, D. 2015. ModelOMatic: Fast and Automated Model Selection between RY, Nucleotide, Amino Acid, and Codon Substitution Models. Systematic biology, 64(1): 42–55.

Woolfit, M. 2009. Effective population size and the rate and pattern of nucleotide substitutions. Biology letters, 5(3): 417–20.

Yang, Z. 1998. Likelihood ratio tests for detecting positive selection and application to primate lysozyme evolution. Molecular biology and evolution, 15(5): 568–573.

Yang, Z. and Nielsen, R. 1998. Synonymous and nonsynonymous rate variation in nuclear genes of mammals. Journal of Molecular Evolution, 46(4): 409–18.

Yang, Z. and Nielsen, R. 2002. Codon-substitution models for detecting molecular adaptation at individual sites along specific lineages. Molecular biology and evolution, 19(6): 908–917.

Yang, Z. and Nielsen, R. 2008. Mutation-selection models of codon substitution and their use to estimate selective strengths on codon usage. Molecular biology and evolution, 25(3): 568–579.

Yang, Z., Nielsen, R., and Hasegawa, M. 1998. Models of amino acid substitution and applications to mitochondrial protein evolution. Molecular biology and evolution, 15(12): 1600–1611.

Zhang, J. 2000. Rates of conservative and radical nonsynonymous nucleotide substitutions in mammalian nuclear genes. Journal of Molecular Evolution, 50(1): 56–68.

